# Community driven dynamics of oscillatory network responses to threat

**DOI:** 10.1101/652834

**Authors:** M Muthuraman, VC Chirumamilla, N Koirala, AR Anwar, O Tüscher, J Vogt, P Horstmann, B Meyer, GA Bonano, G Gonzalez-Escamilla, S Groppa

## Abstract

Physiological responses to threat stimuli involve neural synchronized oscillations in cerebral networks with distinct organization properties. Community architecture within these networks and its dynamic adaptation could play a critical role in achieving optimal physiological responses.

Here we applied dynamic network analyses to address the early phases of threat processing at the millisecond level, describing multi-frequency (theta and alpha) integration and basic reorganization properties (flexibility and clustering) that drive physiological responses. We quantified cortical and subcortical network interactions and captured illustrative reconfigurations using community allegiance as essential fingerprints of large-scale adaptation.

A theta band driven community reorganization of key anatomical regions forming the threat network (TN) along with transitions of nodes from the dorsal attention (DAN) and salience (SN) circuits predict the optimal physiological response to threat. We show that increase flexibility of the community network architecture drives the physiological responses during instructed threat processing. Nodal switches modulate the directionality of information flows in the involved circuits.

These results provide a captivating perspective of flexible network responses to threat and shed new light on basic physiological principles relevant for the development of stress- and threat-related mental disorders.

---

Human responses to threat and stress necessitate highly adaptive but orchestrated responses in the involved brain circuits. Brain oscillations, which play a key role in the coordination of large-scale brain networks, drive physiological responses to threat, and their phase switches determine excitability states of the involved cerebral nodes ^1^. Rapid, temporary shifts of excitability upon stressors involve orchestrated interactions between core components of the threat circuitry, namely the dorsomedial prefrontal cortex (dmPFC), hippocampus and amygdala. These regions could influence primarily adaptive characteristic responses of brain circuits to threat, facilitating coping behavior ^2^. Addressing oscillatory activity in brain networks could delineate real-time processing and unmask involved network nodes illustratively describing the in-phase synchrony of brain oscillations, their states and relation to excitability regulation ^3^.

The core network regions coordinating threat-processing influence the activity in the interconnected areas. Furthermore, the limbic regions of this threat circuit form part of networks mediating the detection and integration of behaviorally relevant stimuli, for instance the salience network (SN) ^4^. Since the involved networks are highly variable and exhibit spontaneous but also task related dynamics ^5^. Identification of the network properties related to threat processing and the specific interrelation among involved brain regions at different networks are of essential importance for understanding mental health related to physiological threat processing, while abnormalities in network associations could be directly and causally linked to mental disorders.

Our conceptual framework is motivated by recent theoretical advances in network science ^6^. These approaches can characterize complex systems as brain responses by delineating components and mapping interactions between interconnected regions ^7^. Addressing the dynamic modification of the networks at different scales allows us to deduce how information is processed between the nodes in the network. Resting state fMRI has been repeatedly used to characterize network dynamics during threat processing ^8^. However, fMRI has, low temporal resolution in seconds, while brain oscillations occur in the millisecond range. Moreover, distinct processes possess a frequency specificity of their evoked responses that cannot be captured by fMRI. Therefore, Electroencephalography (EEG) is ideal to address oscillatory activity at the optimal temporal scale and could help us in the link with modern network science brain responses^9,10^.

Previous evidence suggests that in rodents, oscillations at the theta range (4–8 Hz) support amygdala-prefrontal coordination and contribute to physiological threat processing^11,12^. In non-human primates, the emergence of theta oscillations supports the synchronization of amygdala-to-prefrontal circuits, that serves as a mechanism for long-range communication and directional information transfer between subcortical and cortical structures during threat processing ^13^. In humans, recent studies have shown prominent theta power increases in relation to threat processing at the prefrontal, frontal and midline channels, whereas alpha activity decrease was observed at the parietal and occipital channels ^14,15^.

To address the network dynamics behind threat induced brain responses, we looked for the community network characteristics that are described as properties of functionally specialized sub-networks. The latter are defined as groups of highly inter-connected nodes which have very few connections to nodes in different groups ^16,17^. To capture the dynamic network information processing and behavior in these community networks, we combine several advanced computational algorithms. These include measures of clustering behavior, that capture the capacity to form triangular interconnected communities and flexibility measurements that mirror the extent to which these regions change their community allegiance over time. These measure can effectively tract network reorganization and efficiently quantify the ability to reconfigure to different task demands ^18^.

In the current study, we use an instructed threat paradigm in which the conditioned stimulus (CS+) is paired with an aversive unconditioned stimulus (US) to examine neural processing during threat ^19,20^. Recent studies indicate that physiological responses to threat processing are depicted as increased cortical excitability at time intervals around 1000 milliseconds after the stimuli presentation. For this study, we selected two time points for the application of Transcranial magnetic stimulation (TMS) pulses: one before initiation of threat processing and one at a physiologically relevant time-window. In order to look at the causal network dynamics over a short temporal scale, we take advantage of the EEG ideal temporal resolution and use a non-linear state space modelling approach, which uses dual extended Kalman filtering (DEKF), in a method known as temporal partial directed coherence (TPDC) ^21,22^.

We hypothesized that looking into simultaneous EEG-TMS data, while modulating threat processing through dmPFC stimulation at distinct time intervals, we could uncover local and global network changes at specific neuronal circuits, namely the DAN, SN and FN. Characterization of network reorganization at specific oscillations should also unveil drastic community changes, redefining information flows between nodes. In this line, we evaluate whether network re-configuration is depicted as a change in the region’s assignment to a specific community, or whether network community structure directs the directionality of information flows in the involved circuits.

## Materials and Methods

### Subjects

45 healthy subjects (22 female, mean age 28 ± 5.48 years) were included in our study. The study protocol was approved by local ethics committee (Medical faculty, Johannes Gutenberg University of Mainz) and informed written consent was taken from all participants before the beginning of experiment. All the participants had two visits to the lab, in which during the first visit MRI data was acquired. During the second visit, an instructed threat paradigm (in experiment 1) or an instructed threat paradigm with TMS (experiment 2) was performed.

### MRI data acquisition

Images were acquired using a 3 Tesla MRI scanner (Magnetom Tim Trio, Siemens Healthcare, Erlangen, Germany) equipped with 32-channel head coil at the Neuroimaging Center (NIC) Mainz, Germany. A Magnetization-prepared rapid gradient-echo (MP-RAGE) sequence (Repetition Time [TR] = 1900ms; Echo Time [TE] = 2.54ms; Inversion Time [IT] = 900; Pixel Bandwidth = 180; Acquisition Matrix = 320, 320; Flip Angle = 9°; voxel size = 0.8125, 0.8125 mm; Slice Thickness = 0.8 mm) was used.

### Experiment 1 (instructed threat task)

First, subjects (N=19, 11 female, mean age 27.4 ± 4.32 years) were asked to sit in a chair and painful electric stimuli were applied to the dorsal part of left hand using a surface electrode connected to a DS7A electrical stimulator (Digitimer). Individual pain ratings on a scale from 0 (no pain) to 10 (intense pain) were acquired. An intensity representing a pain level of 7 was used during the experiments. The instructed threat task was developed using the Cogent toolbox (http://www.vislab.ucl.ac.uk/cogent_2000.php) in Matlab 2006B (MathWorks). The task consisted of presenting two visual stimuli: circle and square **(Fig 1 A)**; and a fixation cross during the inter-trial interval (ITI) **(Fig 1 B)**. Participants were instructed that the circle stimulus (CS+) is associated with the electric shock (UCS) with a probability of 33% randomized between (1-5 seconds) at the time the stimulus is presented on screen; while stimulus square (CS−) is not associated with any threat. Stimuli were presented pseudo-randomly on screen for 5 sec and the ITI was jittered between 4 and 6 sec. The paradigm consisted of 60 stimuli (36 CS+, 24 CS−). High-density EEG was recorded from 256 channels throughout the experiment (Net Station 5.0, EGI, USA). Electrode impedances were kept under 50 KΩ during the whole experiment. A sampling frequency of 250 Hz was used. This study was divided into 3 sessions, where each session lasted for around 5 minutes and 3 min breaks were provided in between sessions. After each session the level of experienced threat was rated in a scale from 1 to 10 by each participant with a questionnaire.

**Figure 1:**
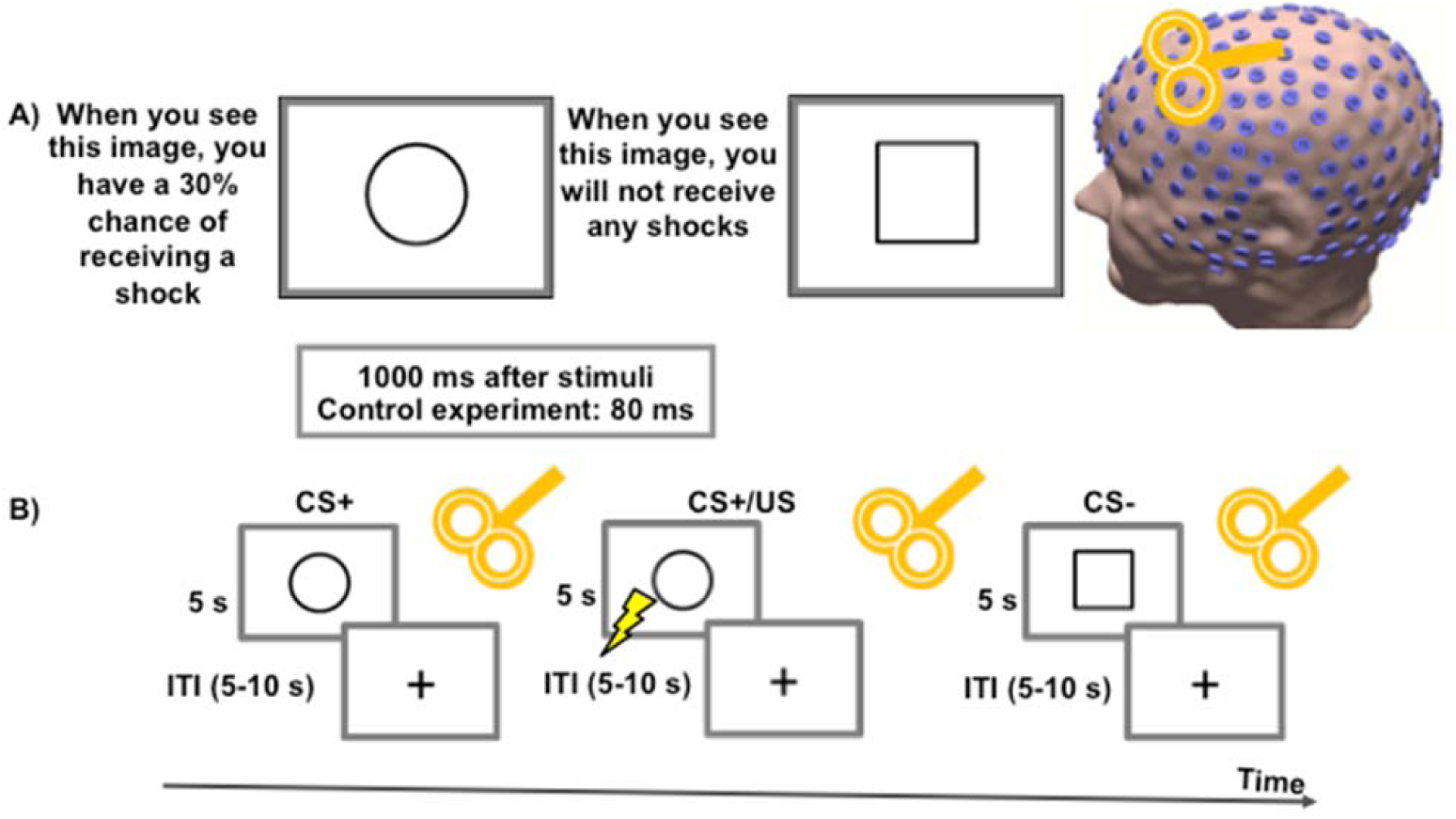
The schematic figure for the instructed threat paradigm used in this study. A) Shows the symbols which were used, when the circle was presented, there was a 30% probability of shock, whereas the square had no probability of shocks. B) Gives an example of the temporal scale of the stimuli presentation, each symbol was presented for 5 seconds, and after either 80 ms or 1 second a neuronavigated single pulse TMS was applied to the right dorsomedial prefrontal cortex (dmPFC). Followed by a fixation cross with an inter-stimulus interval of (5-10) seconds and continued with either CS+ or CS− stimuli in a random manner.

### Experiment 2 (instructed threat task with concomitant dmPFC-TMS)

To evaluate the involvement of the dmPFC in threat processing and network dynamics, a second experiment (Experiment 2) was conducted in another cohort of healthy subjects (N=26, 11 female, mean age 28.6 ± 6.64 years) that includes the instructed threat task as in experiment 1 together with application of single TMS pulses over the right dmPFC, after 1 sec from stimulus onset. The MNI (Montreal Neurological Institute) coordinate for the dmPFC ([10 12 58]) was obtained from a previous fMRI study (1). Individual coordinates were determined using the corresponding MRI in SPM8 (http://www.fil.ion.ucl.ac.uk/spm). Location of TMS pulses delivery, coil position and orientation were controlled throughout the experiment using a neuronavigation system (Localite TMS navigator, Germany). At the stimulation site the TMS coil was placed tangentially to the scalp surface and oriented in a medial to lateral position at a 45° angle away from the midline with the handle pointing backwards. TMS pulses were applied in biphasic pulse configuration using a figure-8 coil connected to Magstim Rapid^2^ (Magstim, UK). The intensity of TMS pulses was set to 110% RMT (resting motor threshold). RMT was calculated as the minimum stimulus intensity required eliciting motor evoked potentials of amplitude 50 *μV* in 5 out of 10 consecutive trials at rest ^23^. The paradigm consisted of 90 stimuli (54 CS+, 36 CS−). The condition specific (CS−: no threat, CS+: threat) trials were considered and the trials in which shock was applied were removed from the analyses. We repeated the experiment 2 but applying TMS at 80ms as a control experiment for TMS modulation on the network dynamics **(Fig 1 B)**. Subjective threat ratings were also acquired in experiment 2.

### EEG data preprocessing

EEG data preprocessing and part of the spatial filter analyses were analyzed using MATLAB2015a and the fieldtrip toolbox ^24^. The data used in the initial preprocessing steps were blind to researcher. Initially, EEG data was re-referenced to the common grand average reference of all EEG channels and epoched from –2.0 to 4.0 s (0 - being the visual stimuli). These epoch trials were used for the purpose of filtering only, for all subsequent analyses the time interval for the epochs was –0.25 to 1.5 s. The preprocessing pipeline was adapted from the Fieldtrip toolbox explained detail in ^19^. For experiment 1, the EEG data was directly subjected to independent component analyses (FastICA) to remove the components representing the muscle artifacts, eye blinks, eye movements and line noise. For experiment 2, TMS-EEG data, a period of –5 to 20 ms relative to the TMS pulse was first cut out and excluded to remove the ringing artifact. The pre-ringing and post-ringing epochs were subject to independent component analysis (FastICA) to remove the components representing the exponential decay artifact, residual muscle artifacts, eye blinks, eye movements, line noise and other muscle artifacts unrelated to TMS. On average for the experiment 1, 30 of 256 components (30 ± 4.6, mean ± SD) were rejected, 10-11 were related to the eye artifacts (11 ± 2.68), 5-6 related to line noise (5 ± 2.34) and 12-13 were related to muscle artifacts (12 ± 1.24). For the experiment 2, 36 of 256 components (36 ± 2.3, mean ± SD) were rejected, 2-3 were related to the exponential decay (2 ± 0.74), 4-5 related to line noise (4 ± 1.98), 13-14 were related to muscle artifacts (13 ± 1.16) and 13-14 were related to the eye artifacts (13 ± 1.04). The residual muscle artifacts were visually inspected, removed and interpolated with the cubic interpolation method. A fourth-order Butterworth low-pass filter with a cut-off frequency of 200 Hz was applied to avoid aliasing, which was followed by a band pass filtered between 3 and 45 Hz.

## Heart rate estimation

The heart rate estimation was extracted from the EEG signals using the extended version of the independent component analysis (ICA) algorithm, based on information maximization ^14^ as previously reported ^9^. In the EEG signals, volume conduction is thought to be linear and instantaneous, and it is expected that the sources of cardiac signals are not generally time locked to the sources of EEG activity, reflecting the activity of cortical neurons ^25^. The lCA can accurately identify the time courses of activation and scalp topographies of relatively large and temporally-independent sources from simulated scalp recordings, even in the presence of a large number of low-level and temporally-independent source activities ^26^.

For heart rate detection analysis, the rows of the input matrix *y* are the EEG signals recorded at the 256 electrodes, the rows of the output data matrix *v* = *X y* are time courses of activation of the lCA components, and the columns of the inverse matrix, *X* ^−1^, give the projection strengths of the respective components onto the scalp sensors.

In general, and unlike PCA, the component time courses of activation will be non-orthogonal. Corrected EEG signals can then be derived as *y*′= (*X*)^−1^*v*′, where *v*′ is the matrix of activation waveforms, *v*, with rows representing sources of cardiac artifacts which are then extracted for further estimations from each participant. In total for experiment 1 we concatenated the 36 CS+ trials to take a total of 180 seconds and 24 CS−trials to take 120 seconds. For experiment 2 we concatenated the 54 CS+ trials to take 270 seconds and 36 CS−trials to take 180 seconds.

### Reliability check of the EEG signals using inter-trial phase coherence (ITPC) analyses

Single trial data were first decomposed into their time-frequency representation by using the multitaper method ^27^. In this method the spectrum is estimated by multiplying the data with *K* different windows (i.e. tapers). In this study, *K* = 7 orthogonal tapers were used with good leakage and spectral properties, the discrete prolate spheroidal sequences (DPSS) are applied ^28^. ITPC were computed using the Fieldtrip toolbox^24^. ITPC reflects the consistency of the phase values across trials, at different frequencies, and at single electrodes. The electrodes were grouped in line with the five lobes to have a global robust measure over a certain lobe instead of selecting certain electrodes. The ITPC was estimated in two frequency bands separately namely theta (4-7 Hz) and alpha (8-13 Hz). This measure is stimulus-locked and independent of amplitude changes. A value of 0 represents absence of synchronization and a value of 1 indicates perfect synchronization. The baseline activity was taken as a reference, and was calculated as the average at each frequency band and across conditions, from –250 to 0 ms before visual stimuli. The change with respect to the baseline interval at 6 windows of 250 ms each after the visual stimulus was then extrapolated up to 1500 ms. Finally, the ITPC difference between the CS+ and CS− was estimated. The significance levels of the ITPC are assessed using surrogate data by randomly shuffling 1000 times the single-trial spectral estimates from different latency windows during the baseline period.

### Reconstruction of brain activity

The forward problem is the computation of the scalp potentials for a set of neural current sources. An established procedure was used by estimating the lead-field matrix with specified models for the brain; a volume conduction model with a finite-element method (FEM) is used ^29^. For the forward modelling the surfaces of the compartments like the skin, skull, csf, gray, and white matter extracted from the individual T1 MRI, and individual electrode locations were used. The forward modeling and the source analysis were done in FieldTrip ^24^. The lead-field matrix (LFM) contains information about the geometry and conductivity of the model. The complete description of the solution for the forward problem has been described previously ^30^. A full description of the beamformer linear constrained minimum variance spatial filter is given elsewhere ^31^. The output of the beamformer at a voxel in the brain can be defined as a weighted sum of the output of all EEG channels. The weights determine the spatial filtering characteristics of the beamformer and are selected to increase the sensitivity to signals from a voxel and reduce the contributions of signals from (noise) sources at different locations. The frequency components and their linear interaction are represented as a cross-spectral density (CSD) matrix. In order to visualize power at a given frequency range, a linear transformation was used based on a constrained optimization problem, which acts as a spatial filter ^32^. The spatial filter assigned a specific value of power to each voxel. For a given source the beamformer weights for a location of interest are determined by the data covariance matrix and the LFM. A voxel size of 5 mm was used in this study, resulting in 6676 voxels covering the entire brain. The created source model was then interpolated on the brain regions defined according to the Automatic Anatomic Labeling Atlas (AAL) 90 cortical regions of interest (ROIs) defined in the MNI space ^33^. For each frequency band (theta and alpha) the activated voxels were selected by a within-subject surrogate analysis to define the significance level, which was then used to identify voxels in the regions as activated voxels. Once the brain region voxels were identified, their activity was extracted from the source space. In a further analysis, all the original source signals for each AAL region with several activated voxels were combined by estimating the second order spectra and employing a weighting scheme depending on the analyzed frequency range to form a pooled source signal estimate for each region as previously described separately for both stimulus (Cs+, CS−) ^34^. Finally, the time series difference between the two conditions (CS+, CS−) was obtained for all the following analyses.

### Evaluating dynamic re-organization of the brain networks

Based on the reconstructed brain activity, individual weighted connectivity matrices were obtained for theta and alpha power separately, and for the same 90 regions of interest defined in the AAL atlas. The links or entries in the connectivity matrix represent the theta or alpha power that is in each ROI(*j*) to all other ROIs(*i*). The weighted connectivity matrices were then characterized using various network measures (see below) as implemented in the brain connectivity toolbox ^35,36^ and the dynamics graph metrics toolbox ^37^.

The communities were identified using the Louvain modularity algorithm ^38^ in each individual subject connectivity matrix. To test the robustness of the detected community association at the baseline interval (−250 to 0 ms), we performed 5000 iterations with the Louvain algorithm where the assignment of each region to a particular community was based on the maximum number of times/iteration a region was assigned to a community ^39^. During this process γ, which is the resolution parameter, was varied from (1 to 2.5) in steps of 0.05 to identify a stable γ value to use in the further time intervals. This procedure was repeated for all the other following intervals from (0 ms to 1500 ms) each within a 250 ms window after the visual stimulus (T1 to T6), with a stable (γ =1.65) to detect the different communities.

### Measures of community efficiency

We assessed three network measures for each formed community: flexibility, clustering coefficient and local efficiency. These measures characterize the efficiency of information transfer at different levels (global and local). To measure changes in the composition of communities ^36^, the flexibility of a node is defined to be the number of times that a node changed community assignment throughout the entire session, normalized by the total number of changes that were possible (i.e., by the number of consecutive pairs of layers in the multilayer framework). We then defined the flexibility of the entire community as the mean flexibility over the nodes in that particular community. The clustering coefficient (C) is a parameter of local organization ^40^ reflecting the number of connections between directly neighboring nodes (the topological motif of a triangle), with sparsely interconnected regions showing lower values. The efficiency of a network primarily reflects how information is exchanged between the regions. Local efficiency ^41^ quantifies a network’s resistance to failure on a small scale and is defined as the inverse of the length of the shortest path in the node. For the three network parameters, twenty density intervals (range 0.1–0.6) were calculated, over which we estimated the mean and standard deviation. The range was chosen in such a way that the network was fully connected at the minimum value and fully disconnected at the maximum value ^42^.

### Investigating causal relationships between network nodes

The effective connectivity analysis was performed on all the nodes separately, for each of the three new communities and each frequency separately. Using time-frequency causality, is possible not only to focus on a particular frequency, but also to analyze the dynamics of the causality at that frequency. The time-frequency causality estimation using the TPDC is based on dual extended Kalman filtering (DEKF) ^43^, and allows time-dependent auto regressive (AR) coefficients to be estimated. One EKF estimates the states and feeds this information to the other; the second EKF estimates the model parameters and shares this information with the first. By using two Kalman filters working in parallel with one another, we can estimate both states and model parameters of the system at each time instant. After estimating the time-dependent multivariate (MVAR) coefficients, the next step is to use those coefficients for the calculation of causality between the time series. By calculating the time-dependent MVAR coefficients at each time point, we can also calculate partial directed coherence (PDC) at each time point. The frequency bands taken into account were the theta and alpha. After estimating the TPDC values the significance level was calculated from the applied data using a bootstrapping method ^44^. In short, we divide the original time series into smaller non-overlapping windows and randomly shuffle the order of these windows to create a new time series. The MVAR model is fitted to the shuffled time series and TPDC is estimated. The bootstrapping is performed 1000 times and the average TPDC value is taken as the significance threshold for all our connections. This process is performed separately for each participant. The resulting value is the significance threshold value for all our connections. This process is performed separately for each subject. In this study the open source Matlab package autoregressive fit (ARFIT)^45^ was used for estimating the autoregressive coefficients from the spatially filtered source signals of the identified nodes in the three new communities. We applied time reversal technique TRT ^46^ as a second significance test on the connections already identified by TPDC using a data-driven bootstrapping surrogate significance test. We have previously applied this type of non-linear time-frequency causality both in EEG ^22,31^ and functional modality datasets ^21^.

## Statistical analyses

We checked the normality of the data using the Shapiro-Wilk test and the sphericity was checked with Mauchly’s sphericity test. The behavioral (threat) ratings and the heart rate values were tested for significance (p < 0.01) between the two stimuli CS+ and CS− using paired T-test. The three network measures for each of the three networks and two experiments were tested for significance separately using a two-way factorial ANOVA, within subject factor (n=2; conditions, time). The TPDC values between the regional source signals were tested for significance using a paired T-test. The significant differences were tested for each frequency, each community and from each two following time windows separately (for ex: baseline Vs T1; T1 Vs T2; T2 Vs T3; T3 Vs T4; T4 Vs T5; T5 Vs T6).

The Pearson correlation coefficient was estimated between the behavioral ratings (difference between CS+ and CS−) and the heart rate (difference between CS+ and CS−). Finally the network parameters and the effective connectivity values from all the time windows (T1-T6) were correlated separately with the behavioral ratings and the heart rate. The Bonferroni correction was performed for all the post-hoc tests and was considered significant at p < 0.05.

## Results

We began by asking whether regions of the brain change their community attribution at theta and alpha frequency bands derived from high-density EEG during threat processing. We then examined three parameters of network organization (flexibility, clustering and local efficiency). We finally studied whether dynamic network characteristics using effective connectivity measures can be reliably traced during threat processing in healthy subjects, and searched for associations between network dynamics, behavioral and electrophysiological responses during threat processing.

### Behavioral quantification of threat states and heart rate analyses

The behavioral ratings of threat were higher in the threat (CS+) condition when compared to the no-threat (CS−) condition (p < 0.001; **Fig. 2*A***). The estimated heart rate showed clear increases for the CS+ compared to the CS− (p < 0.001; **Fig. 2*B***). Primary behavioral effects from experiment 1 (no TMS) were replicated in experiment 2 (TMS) for both increased ratings to CS+ (p < 0.001; **Fig. 2*A***) and heart rates (p < 0.001; **Fig. 2*B***).

**Figure 2:**
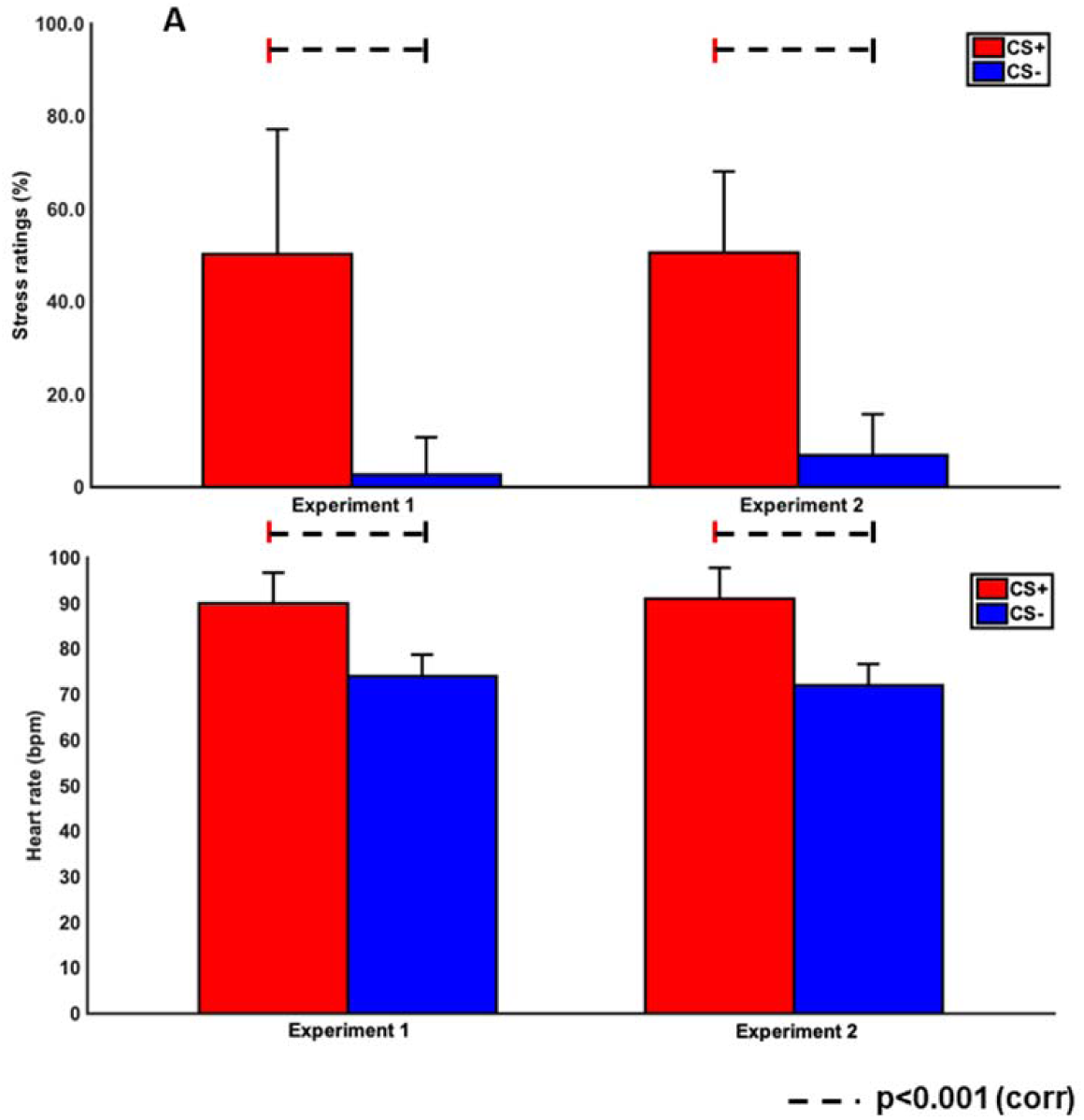
The behavioral stress ratings mean and standard deviation for both the experiments are shown in A) and the heart rate mean and standard deviation for both the experiments are shown in B). The dark grey bar represents the CS+ stimuli and the light grey bar represents the CS− stimuli. The dashed line indicates the significant difference between the two stimuli.

The correlation analyses between heart rate and threat ratings showed significant associations in both experiment 1 (r = 0.56; p = 0.002) and experiment 2 (r = 0.49; p = 0.005).

### Threat processing related inter-trial phase coherence changes

The frontal theta showed a significant inter-trial phase coherence (ITPC) increase in between the baseline (−250-0 ms) and the four subsequent time windows from (0 to 1000 ms) (experiment 1 p < 0.001; experiment 2 p < 0.001; **Fig. 3**). The occipital alpha showed decreased ITPC between the two conditions (CS+ and CS−) but the difference was only significant between the baseline (–250-0 ms) and the subsequent three time windows from (0-750 ms) in both experiment 1 (p < 0.001) and experiment 2 (p < 0.001). The ITPC in the theta frequency showed an inverted U-shape like temporal pattern, increase in the first two windows (0−250 and 250-500 ms) and then reduced in the two subsequent time windows (500-750 and 750-1000 ms) in both experiments. The increase in ITPC of the frontal lobe showed the robust nature of the oscillatory response of each trial for the threat processing stimuli. The occipital alpha showed the opposite behavior, a decrease in the first two windows (0−250 and 250-500 ms) after the visual stimuli followed by an increase in the time windows 500-750 and 750-1000 ms also in both experiments. The decrease of ITPC in the occipital lobe showed the specificity of attention for threatful stimuli.

**Figure 3:**
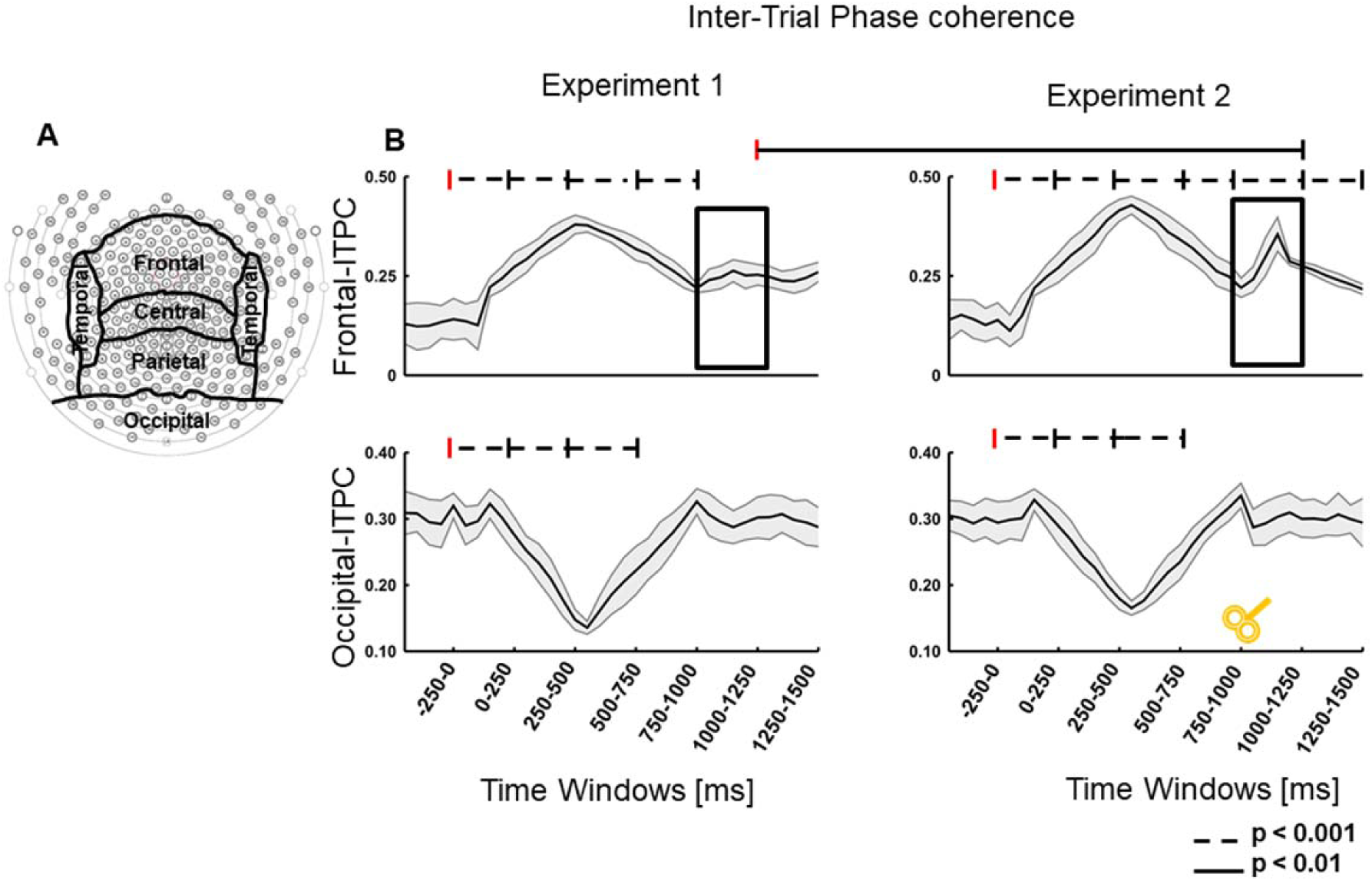
A) Shows the subdivision of the lobes for the estimation of the inter-trial phase coherence. B) First row shows the frontal inter-trial phase coherence (ITPC) for both the experiments and time windows, starting from the baseline (−250 to 0 ms) followed by six time windows (T1-T6) each 250 ms, the second row shows the occipital ITPC. The dashed black line indicates the significant differences and the red vertical line indicates the window to which the comparison was done. The black boxes in B) indicate the change in ITPC between the experiment 1 and experiment 2. The TMS coil indicates the time of application of single pulse TMS.

### TMS induced inter-trial phase coherence changes

In experiment 2 the ITPC at the theta band was perturbed by the TMS in the dmPFC at 1000 ms and induced a significant increase (p < 0.01) in the frontal theta but did not affect the occipital alpha (**Fig. 3**). We were able to modulate the ITPC using TMS at the dMPFC in the frontal lobe during the time interval of threat processing. The control experiment of TMS at 80 ms did not induce any significant change in the frontal lobe indicating choosing the correct temporal window for perturbation is vital.

### Network communities of the source signals

The community detection analyses at both the theta and the alpha bands identified nine modules during the baseline interval (−250 to 0 ms; **Fig. 4**). The configuration of the communities was anatomically differentiated, namely community 1 comprised frontal regions; community 2 included basal ganglia; community 3 and community 7 encompassed fronto-parietal regions. Community 4 and community 5 included parietal and occipital regions respectively; whereas all the temporal regions were incorporated in community 6. Community 8 included sensorimotor regions and community 9 comprised hippocampus and amygdala. When testing the robustness of the community association, this was stable over the 5000 iterations, i.e., the regions were assigned to the exact same community in 80 ± 6% of the iterations.

**Figure 4:**
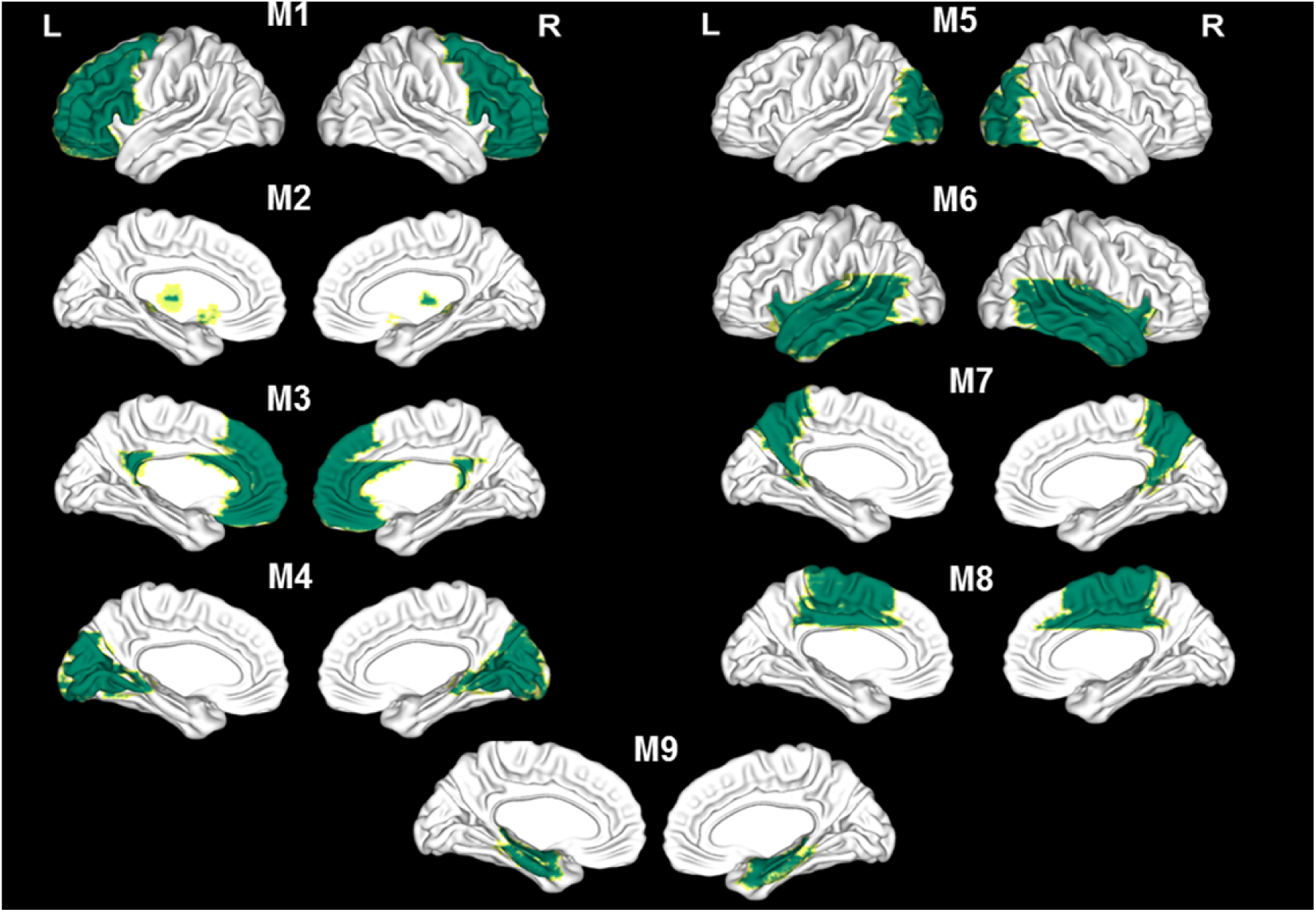
Shows the identified nine communities from (M1 to M9) at the baseline time window (−250 to 0) ms. The community 1 comprise of mainly frontal regions. The community 2 includes basal ganglia; community 3 and community 7 encompassed fronto-parietal regions. Community 4 and community 5 include parietal and occipital regions; community 6 includes temporal regions. Community 9 comprised of hippocampus and amygdala and community 8 included central regions.

After the visual stimuli in the experiment time windows 0 to 1500 ms all the nodes from the community 4 did not change the nodal alliance (Supplementary Table 1). However, in all the other communities either all the nodes as in community 5 or only some nodes altered their alliance to the known communities. The three well-known communities were the dorsal attention (DAN), salience (SN) and threat (TN) networks. The nodes that altered the community are marked in three different colors namely red for DAN, blue for SN and green for FN representing each of these three networks (**Supplementary Table 1**).

### Temporal changes in network organization within communities

In the theta band from the three formed networks, the dorsal attention network (**Fig. 5*A***) showed significantly increased flexibility in the experiment 1, factor condition (F_1,18_ = 22.67, p < 0.001) and factor time (F_6,108_ = 12.24, p < 0.001); and in experiment 2, factor condition (F_1,25_ = 20.45, p < 0.001) and time (F_6,150_ = 14.87, p < 0.001). The post-hoc analyses showed significantly higher flexibility for windows from (0 to 1500 ms) in comparison to the baseline window (−250-0 ms) (p < 0.001 for all windows; **Fig. 5*B***). In experiment 2, the window 1000-1250 ms showed a decrease in flexibility with respect to the window 750-1000 ms (p < 0.001). In the alpha frequency band the regions of the DAN showed no flexibility changes with respect to the baseline (−250-0 ms) in both experiments. A flexibility increase in the window 1000-1250 ms in comparison to 750-1000 ms window was observed (p < 0.001) in experiment 2.

**Figure 5:**
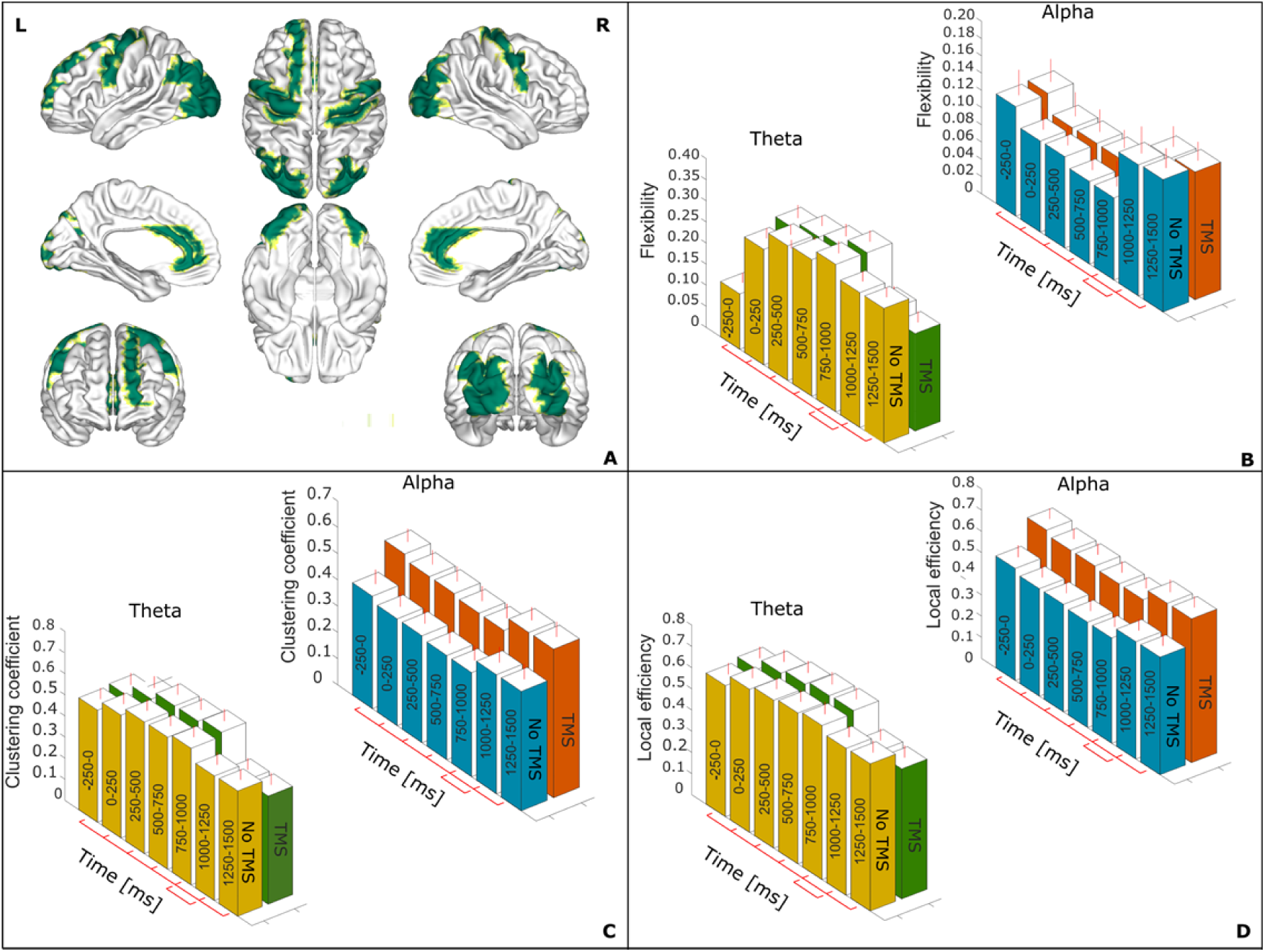
In A) the representative figure with regions comprised in the newly formed dorsal attention network (DAN), the corresponding list is given in Supplementary Table 1. The network parameter flexibility is shown in B) starting from the baseline (−250 to 0) ms window to all the following six time windows (T1-T6) for every 250 ms separately bar plots with mean and standard deviation for theta and alpha frequency bands. C) and D) show the values for the clustering coefficient and the local efficiency. The red line indicates the significant differences between the time intervals all the intervals were compared to the baseline. The interval (750-1000) ms were compared to the (1000-1250) ms.

The clustering coefficient (**Fig. 5*C***) also showed significant increases in the theta frequency band in experiment 1, factor condition (F_1,18_ = 38.74, p < 0.001) and time (F_6,108_ = 19.57, p < 0.001) and experiment 2, factor condition (F_1,25_ = 34.21, p < 0.001) and time (F_(6,150)_ = 17.24, p < 0.001). The post-hoc analyses revealed no significant differences between baseline (−250-0 ms) and all the experiment windows from 0 to 1500 ms (p > 0.05 for all time intervals). The alpha band showed no time changes with respect to the window −250-0 ms in both experiments. An increase of clustering coefficient was detected for the window 1000-1250 ms in comparison to 750-1000 ms, p = 0.006) in experiment 2.

The network local efficiency (**Fig 5*D***), was increased in the theta band in experiment 1, factor condition (F_1,18_ = 42.28, p < 0.001) and time (F_6,108_ = 23.27, p < 0.001) and experiment 2, factor condition (F_1,25_ = 39.65, p < 0.001) and time (F_6,150_ = 19.38, p < 0.001), from −250-0 ms to the 0-1500 ms windows. Only in the experiment 2 (TMS) a significant decrease was observed for the 750-1000 ms)window in comparison to 1000-1250 ms (p < 0.001). In the alpha band the experiment windows (0 to 1500 ms) did not differ from baseline window (–250-0 ms) in both experiments, only an increase for the interval (1000-1250 ms) from (750-1000 ms) was significantly different (p < 0.05) in both experiments.

The regions forming the salience network (**Fig 6A**) showed significant flexibility increases (**Fig 6B**) in the theta band for the experiment 1, factor condition (F_1,18_ = 12.67, p = 0.002) and time (F_6,108_ = 4.24, p = 0.001) and in experiment 2, factor condition (F_1,25_ = 9.24, p = 0.006) and time (F_6,150_ = 5.87, p < 0.001). The post-hoc analyses revealed significant differences between (−250-0 ms) and the windows (0 to 1500 ms) (p < 0.001 for all time intervals) in both experiments. In experiment 2 the interval 1000-1250 ms showed a significant flexibility increase (p < 0.001) with respect to the previous interval 750-1000 ms. In the alpha band a decrease in flexibility was observed in the experiment 1, factor condition (F_1,18_ = 8.48, p = 0.01) and time (F_6,108_ = 3.68, p = 0.009) and in experiment 2, factor condition (F_1,25_ = 5.45, p = 0.03) and time (F_6,150_ = 2.86, p = 0.01). An increase of flexibility was also observed for the interval 1000-1250 ms in comparison to 750-1000 ms, which was only significant in experiment 2 (p < 0.001).

**Figure 6:**
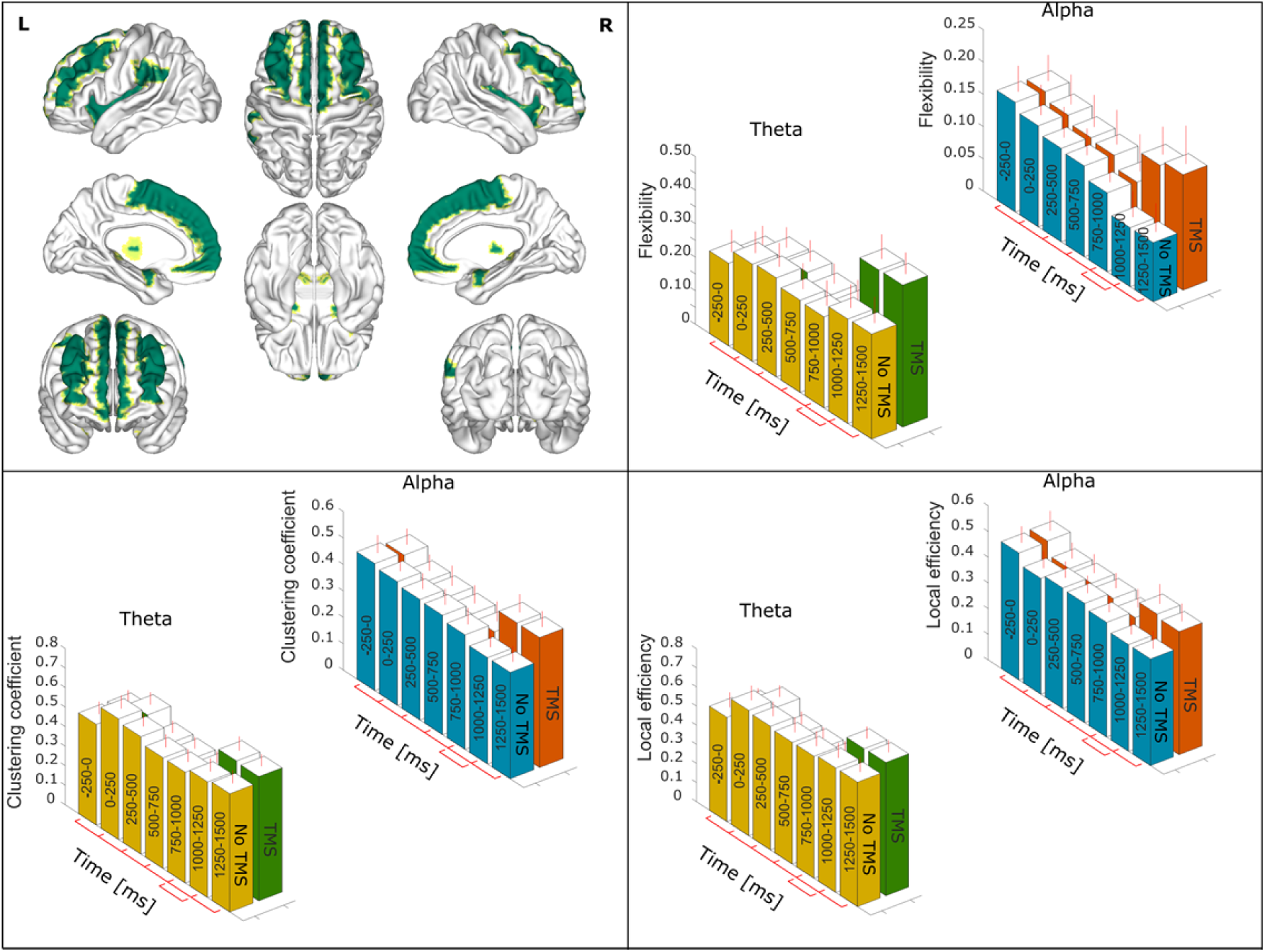
In A) the representative figure with regions comprised in the newly formed Salience network (SN), the corresponding list is given in Supplementary Table 1. The network parameter flexibility is shown in B) starting from the baseline (−250 to 0) ms window to all the following six time windows (T1-T6) for every 250 ms separately bar plots with mean and standard deviation for theta and alpha frequency bands. C) and D) show the values for the clustering coefficient and the local efficiency. The red line indicates the significant differences between the time intervals all the intervals were compared to the baseline. The interval (750-1000) ms were compared to the (1000-1250) ms.

The clustering coefficient (**Fig 6C**) of the theta band also showed significant increases in the experiment 1, factor condition (F_1,18_ = 10.54, p = 0.005) and time (F_6,108_ = 3.64, p = 0.008) and experiment 2, factor condition (F_1,25_ = 8.46, p = 0.009) and time (F_6,150_ = 3.72, p = 0.008). The post-hoc analyses showed clustering increases at all the time intervals from 0 to 1500 ms with respect to the –250-0 ms (p < 0.01 - for all time intervals). In experiment 2 the window 1000-1250 ms showed a significant increase (p < 0.001) in the flexibility with respect to 750-1000 ms. In the alpha band no significant changes in respect to baseline –250-0 ms were detected in experiment 1 or experiment 2. The increase in clustering coefficient for the interval 1000-1250 ms from 750-1000 ms was significant (p < 0.001) only for the experiment 2.

The local efficiency (**Fig 6D**), reflecting network resistance to failure, was increased in the theta band of experiment 1, factor condition (F_1,18_ = 11.25, p = 0.003) and time (F_6,108_ = 4.78, p < 0.001) and experiment 2, factor condition (F_1,25_ = 11.45, p = 0.002) and time (F_6,150_ = 5.46, p < 0.001). Post-hoc analyses revealed differences at all time-windows from 0 to 1500 ms with respect to the –250-0 ms window (p < 0.01 for all time intervals). However, significant local efficiency increases were detected for the interval 750-1000 ms in respect to 1000-1250 ms (p < 0.01) only in the experiment 2 (TMS). The alpha band showed significantly decreased local efficiency in the experiment 1, factor condition (F_1,18_ = 9.24, p = 0.007) and time (F_6,108_ = 2.78, p = 0.01) and experiment 2, factor condition (F_1,25_ = 9.22, p = 0.006) and time (F_6,150_ = 3.18, p = 0.008). The post-hoc analyses showed significant increases in the windows from 0 to 1500 ms in comparison to –250-0 ms (p < 0.01 for all windows).

The regions of threat network **(Fig 7A and B)**, showed significant changes only in the theta band and no significant changes was observed in the alpha band from (−250-0 ms) to the following time windows (0 to 1500 ms). This network showed significantly increased flexibility in the theta frequency of experiment 1, factor condition (F_1,18_ = 15.42, p = 0.001) and time (F_6,108_ = 7.65, p < 0.001) and experiment 2, the factor condition (F_1,25_ = 15.53, p < 0.001) and time (F_6,150_ = 8.96, p < 0.001). Post-hoc analyses revealed higher flexibility in windows (0 to 1500 ms) compared to (−250-0 ms) (p < 0.001 for all windows in both experiments). In experiment 2 the (1000-1250 ms) showed a significant flexibility increases with respect to (750-1000 ms) (p < 0.001).

**Figure 7:**
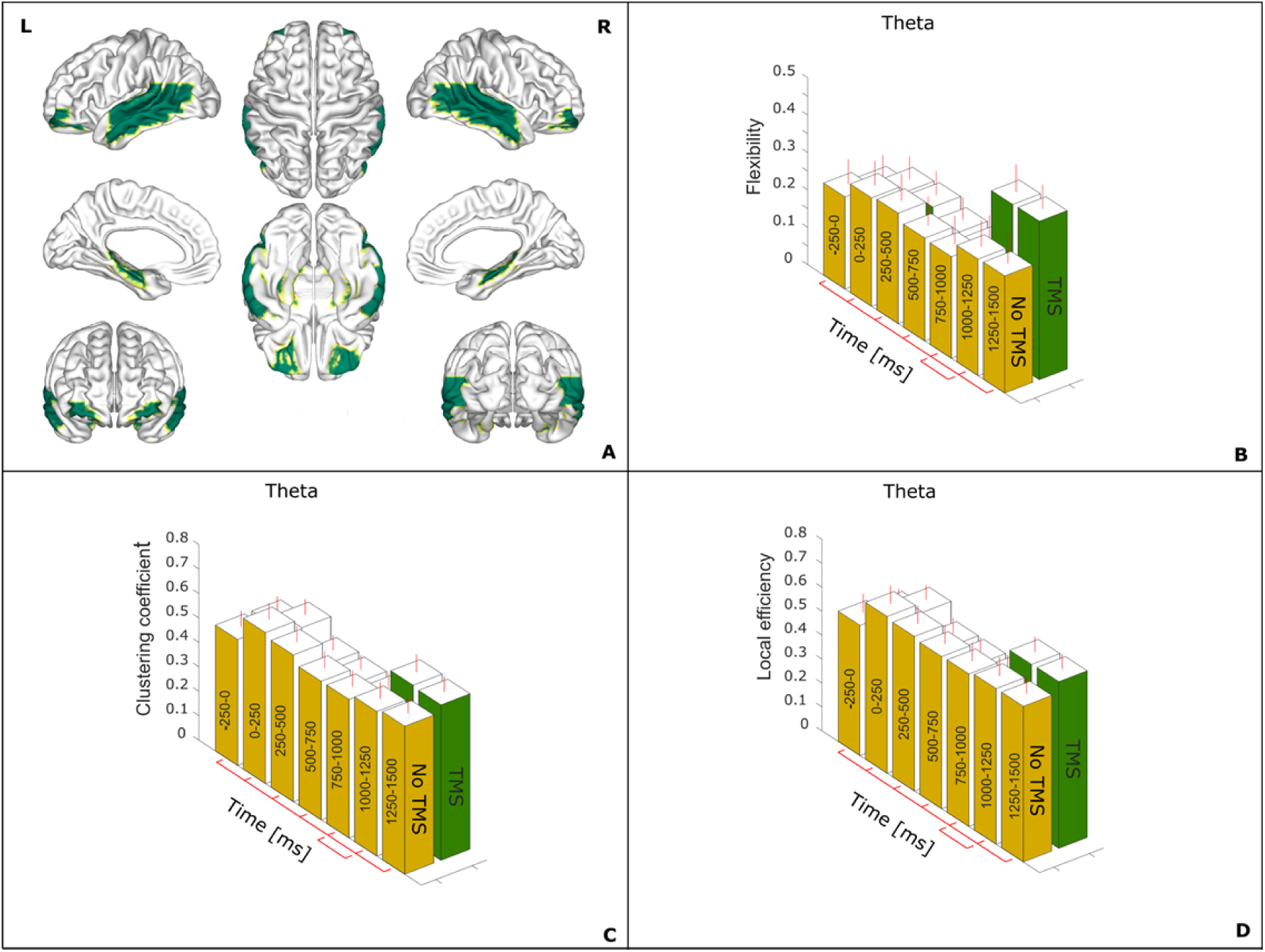
In A) the representative figure with regions comprised in the newly formed threat network (TN), the corresponding list is given in Supplementary Table 1. The network parameter flexibility is shown in B) starting from the baseline (−250 to 0) ms window to all the following six time windows (T1-T6) for every 250 ms separately bar plots with mean and standard deviation for theta and alpha frequency bands. C) and D) show the values for the clustering coefficient and the local efficiency. The red line indicates the significant differences between the time intervals all the intervals were compared to the baseline. The interval (750-1000) ms were compared to the (1000-1250) ms.

The clustering coefficient (**Fig 7C**) also showed a significant theta band increases in the experiment 1, factor condition (F_1,18_ = 16.21, p < 0.001) and time (F_6,108_ = 9.47, p < 0.001) and experiment 2, factor condition (F_1,25_ = 17.24, p < 0.001) and time (F_6,150_ = 7.67, p < 0.001). Post-hoc analyses comparing the (−250-0 ms) and the experiment windows (0 to 1500 ms) showed significant increase (p < 0.001 for all time intervals). In experiment 2 the interval (1000-1250 ms) showed a significant increase (p < 0.001) in clustering coefficient with respect to (750-1000 ms).

The local efficiency (**Fig 7D**) of the theta band was increased in experiment 1, factor condition (F_1,18_ = 12.31, p = 0.001) and time (F_6,108_ = 9.46, p < 0.001) and experiment 2, the factor condition (F_1,25_ = 14.20, p < 0.001) and time (F_6,150_ = 9.35, p < 0.001), followed by post-hoc analyses revealed significant difference from the (−250-0 ms) to the intervals (0 to 1500 ms) (p < 0.001 for all time intervals). However, only in experiment 2 (TMS) a significant increase was detected from (750-1000 ms) to (1000-1250 ms) (p < 0.001). In this community the alpha band did not exhibit any significant differences for both the experiments in none of the analyzed network measures. The network measures flexibility, clustering and local efficiency for the control experiment of single pulse TMS at 80 ms did not change the network connectivity dynamics for all the three communites as shown in Suppl. Fig.1 between the baseline (−250-0 ms) and the first time window (0−250 ms).

### Effective connectivity dynamics

The dynamical, effective connectivity (**Fig 8–10**) analyses were focused only on the difference between the two conditions (CS+ and CS−) and the three newly formed communities. In the baseline time window (−250-0 ms) only information flows that survived surrogate and time reversal technique (p < 0.001) are reported. In **Fig 8 to 10** the information flow at baseline (−250-0 ms) for the three networks is shown: red lines for the theta band and blue lines for the alpha band. Significant changes in the directionality of information flow for experiment time windows in comparison to the baseline are shown in yellow; the thickness of the lines indicates the strength of the information flow.

**Figure 8:**
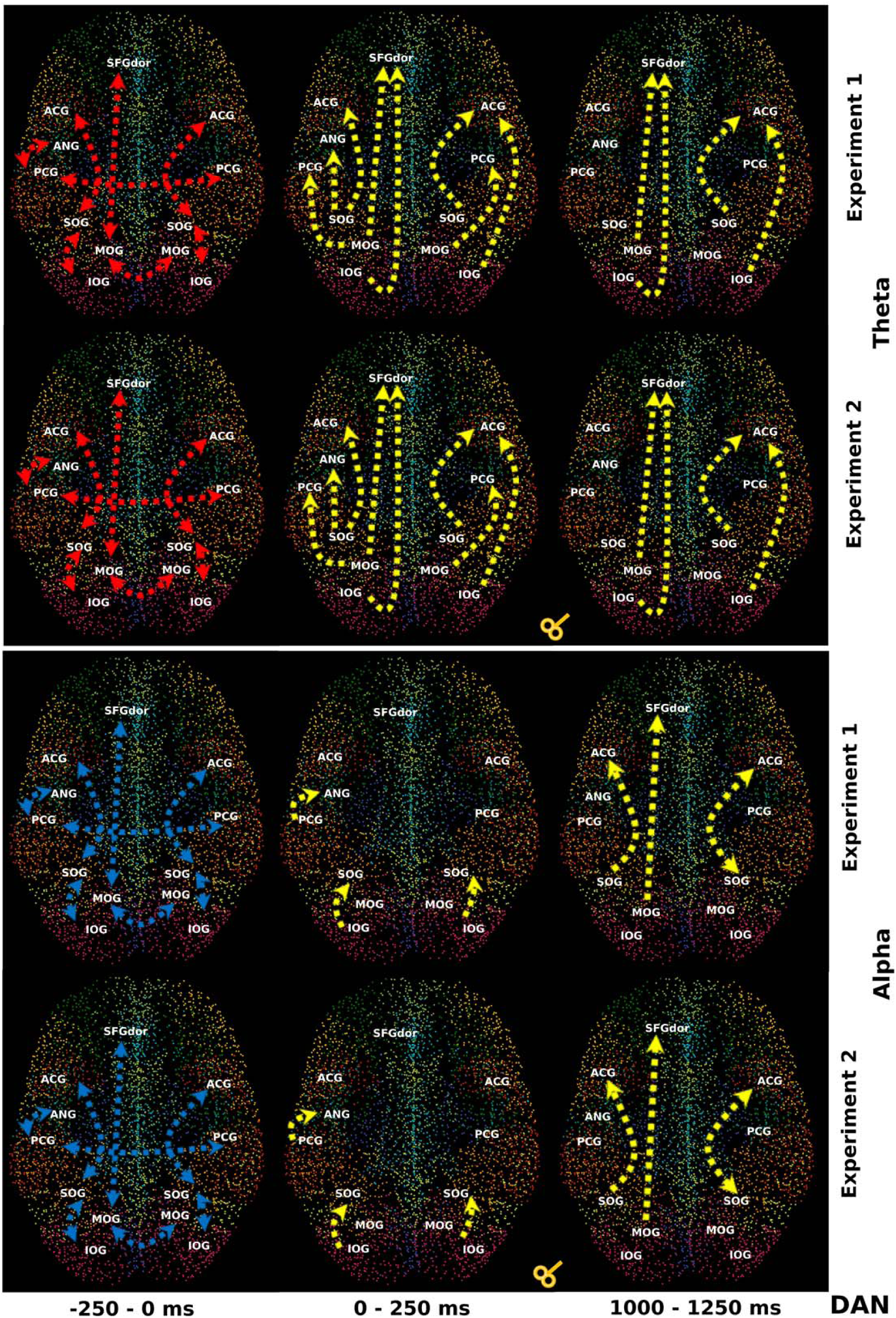
The temporal partial directed coherence (TPDC) information flow between the regions in the dorsal attention network (DAN) are shown in the template brain showing the 9 communities in different colors and the 6676 voxels. The first two rows for the theta and the last two rows for the alpha frequency band separately. The first and third row represents the experiment 1 and the second and fourth row the experiment 2. The red arrows indicate the information flow during the baseline window (−250 to 0) ms and the yellow lines indicates the difference in the information flow to the previous time window and the thickness of the lines indicates the strength of the information flow when compared to the previous time window. *L: Left; R: Right; SFGdor: Superior frontal Gyrus, dorsolateral; ACG: Anterior Cingulate Gyrus; SOG: Superior Occipital Gyrus; MOG: Middle Occipital Gyrus; IOG: Inferior Occipital Gyrus; PCG: Posterior Cingulate Gyrus; ANG: Angular Gyrus;*

**Figure 9:**
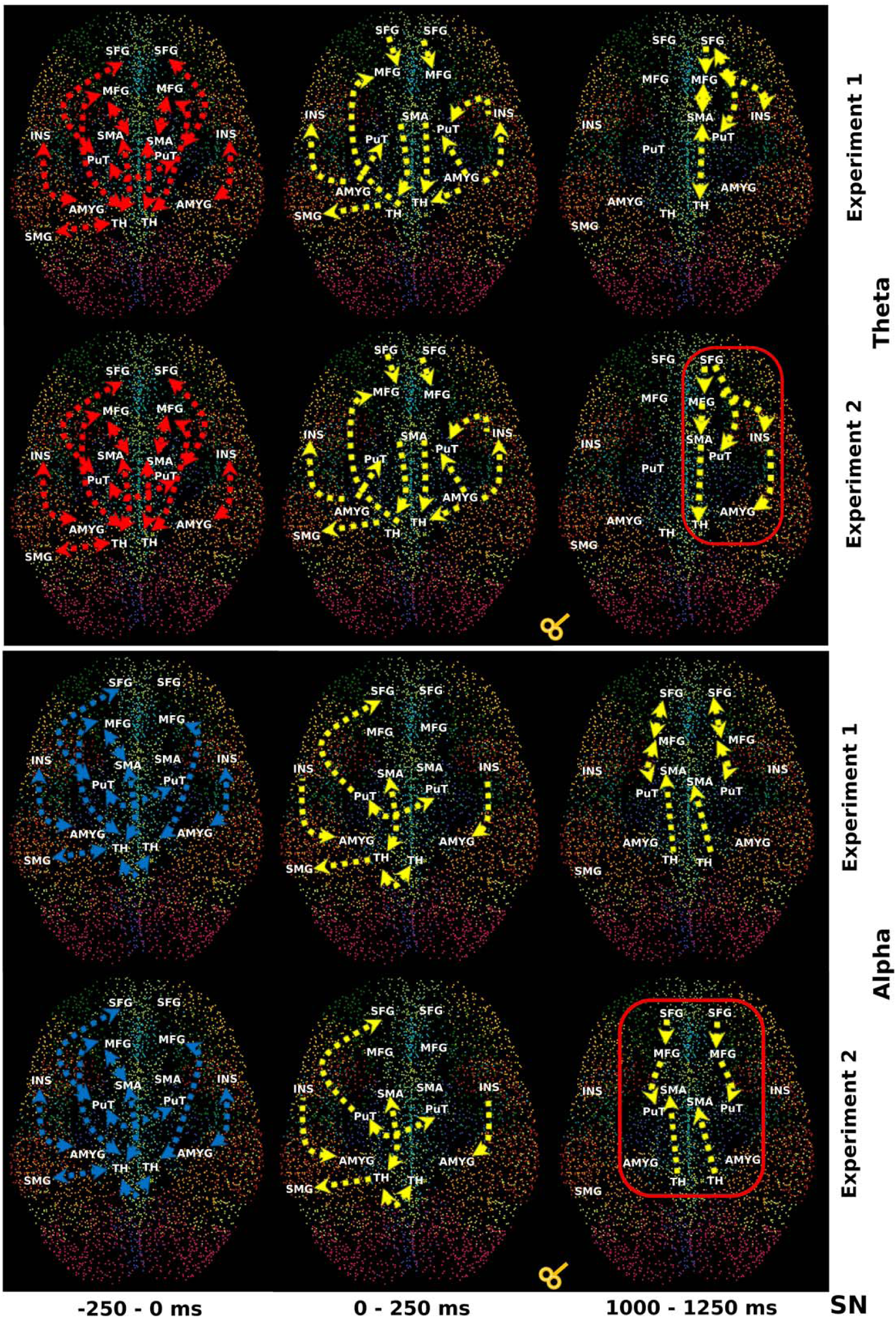
The temporal partial directed coherence (TPDC) information flow between the regions in the salience network (SN) are shown in the template brain showing the 9 communities in different colors and the 6676 voxels. The first two rows for the theta and the last two rows for the alpha frequency band separately. The first and third row represents the experiment 1 and the second and fourth row the experiment 2. The red arrows indicate the information flow during the baseline window (−250 to 0) ms and the yellow lines indicates the difference in the information flow to the previous time window and the thickness of the lines indicates the strength of the information flow when compared to the previous time window. The red box indicates the difference in information flow between experiment and experiment 2. *L: Left; R: Right; MFG: Middle Frontal Gyrus; SMA: Supplementary Motor Area; INS: Insula; AMYG: Amygdala; SMG: Supramarginal Gyrus; PUT: Putamen; THA: Thalamus; SFGmed: Superior Frontal Gyrus, Medial;*

**Figure 10:**
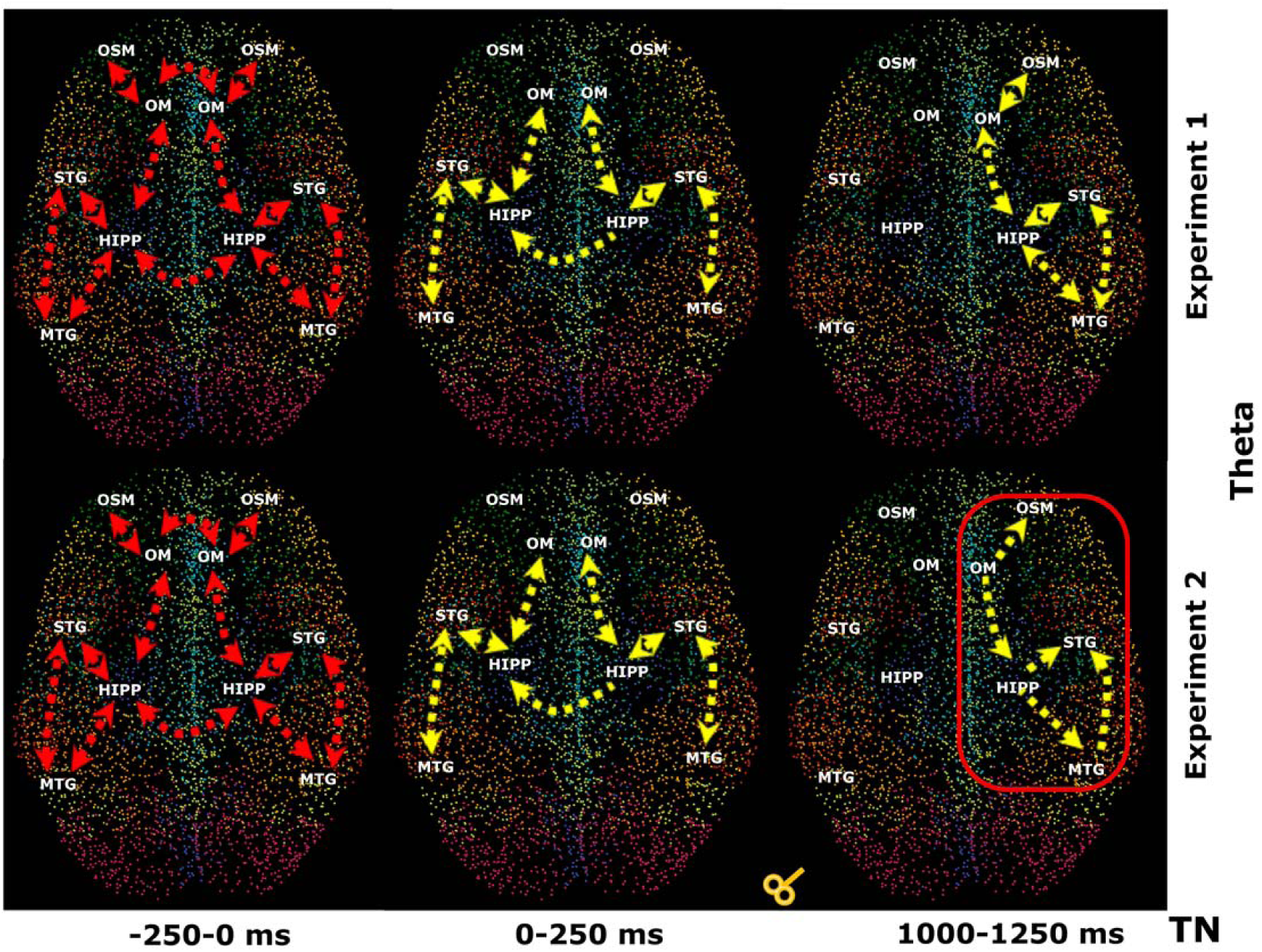
The temporal partial directed coherence (TPDC) information flow between the regions in the threat network (TN) are shown in the template brain showing the 9 communities in different colors and the 6676 voxels. The first two rows for the theta and the last two rows for the alpha frequency band separately. The first and third row represents the experiment 1 and the second and fourth row the experiment 2. The red arrows indicate the information flow during the baseline window (−250 to 0) ms and the yellow lines indicates the difference in the information flow to the previous time window and the thickness of the lines indicates the strength of the information flow when compared to the previous time window. The red box indicates the difference in information flow between experiment and experiment 2. *L: Left; R: Right; HIPP: Hippocampus; STG: Superior Temporal Gyrus; MTG: Middle Temporal Gyrus; ORBsupmed: Superior Frontal Gyrus, medial orbital; ORBmid: Middle Frontal Gyrus, orbital part;*

In the dorsal attention network (**Fig 8**), the information flow of the baseline window (- 250-0 ms) was similar in both experiments and both frequency bands (theta, alpha). In this window the information flow was bi-directional and largely restricted to intra-hemispheric connections with only two inter-hemispheric connections. In the theta band the connections at (0−250 ms) (immediate after visual stimuli) changed from bi-directional to uni-directional. The information flows in this band were restricted to intra-hemispheric connections from occipital to frontal regions in both experiments. At (1000-1250 ms) fewer connections showed changes but they had increased strength flow (thicker yellow lines in figure 8), specifically MOG (left) to SFGdor (left), IOG (left) to SFGdor (left), SOG (right) to ACG (right) and IOG (right) to ACG (right). For the alpha band, at (0−250 ms) uni-directional connections remain between IOG and SOG in both hemispheres, and PCG (left) to ANG (left). At the interval (1000-1250 ms) three new connections had increased strength with respect to (750-1000 ms); SOG (left) to ACG (left), MOG (left) to SFGdor (left) and SOG (right) to ACG (right). In the salience network (**Fig 9**), at baseline (−250-0 ms) of the theta band only bi-directional connections were observed between the regions; most were intra-hemispheric except between left and right putamen, in both experiments. At (0−250 ms) the information flow turned uni-directional and intra-hemispheric, with increased strength flow with respect to baseline (−250-0 ms). In the experiment 1 all remained connections at (1000-1250 ms) were restricted to the right-hemisphere and exhibited bi-directional information flow, except from the insula to amygdala. TMS modulation of the theta band was visible in the experiment 2 at (1000-1250 ms), where the existing connections turned uni-directional. Connections were from frontal cortex to the supplementary motor area (SMA) and sub-cortical regions (putamen and thalamus) and from insula to amygdala.

In the alpha band, at baseline (−250-0 ms) the information flow of the salience network was similar to the theta band, except for an inter-hemispheric connection between the bilateral thalami. At (0−250 ms) the connection strength was higher in comparison to the baseline window (−250-0 ms) and only three additional bi-directional connections remained, whereas four connections turned uni-directional. At (1000-1250 ms) connections remained in both hemispheres; all were intra-hemispheric and bi-directional, except from thalamus to the SMA in case of the experiment 1. In experiment 2 remained connections turned uni-directional, these were from SFG to MFG, MFG to putamen and thalamus to SMA in both hemispheres.

In the threat network (**Fig 10**) the theta frequency connectivity at baseline (−250-0 ms) showed again only bi-directional connections between the regions; most of them were intra-hemispheric except between bilateral ORBmid and HIPP, in both experiments. At (0−250 ms) the information flow of most connections remained bi-directional and intra-hemispheric, except between bilateral hippocampi, which turned uni-directional. The strength of the connections was significantly increased with respect to baseline window (−250-0 ms). At (1000-1250 ms), in the experiment 1 all connections remained bi-directional and were restricted to the right hemisphere. Theta band TMS modulation was visible in the experiment 2, where all the connections turned uni-directional. The connection strength was higher for the existing connections, except for ORBmid (right) to ORBsupmed (right), which was weaker at this time window.

### Correlation between electrophysiological and behavioral indicators of threat processing

Only the significant correlations that survived Bonferroni correction and were attested in both experiments are reported. We found a significant correlation between the frontal theta ITPC positive slope (−250-0 to 250-500 ms) and the heart rate in both experiment 1 (r = 0.70; p = 0.014) and experiment 2 (r = 0.61; p = 0.012). A significant correlation was also found between the slope of the windows (−250-0 to 0−250 ms) for the network flexibility parameter in the theta band of CS+ stimuli and heart rate in the salience and threat networks, in both experiment 1 (r = 0.68; p = 0.003; r = 0.56; p = 0.005) and experiment 2 (r = 0.58; p = 0.005; r = 0.61; p = 0.004). The correlation between the effective connectivity from the time interval (0−250 ms) and the difference in threat ratings were only significant for the salience network; specifically for two connections: INS (left) to AMGY (left) (r = 0.40; p = 0.006) and TH (left) to SMA (left) (r = 0.37; p = 0.009). The results from correlation analyses between the theta band (0−250 ms) window effective connectivity and the heart rate are listed in **Supplementary Table 2**, for each community and for both experiments. Correlation results between the alpha band (0−250 ms) window effective connectivity and the heart rate are listed in **Supplementary Table 3**, for each community and for both experiments. The correlation results between the theta band (1000-1250 ms) window effective connectivity and the heart rate are listed in **Supplementary Table 4**, separately for each community and experiments.

## Discussion

Decades of research has attempted to decipher the neurobiological basis of threat processing. In the current study, we demonstrate that a quick shift of cognitive, associative learning and classical conditional mechanisms, mirrored by particular network crosstalk, is needed in order to shape our neurobiological response to aversive stimuli. Up until now, methodological difficulties in quantifying the neural elements of the processing network at high temporal resolution meant that it was not possible to arrive at a personalized non-invasive description of human behavior to threat. The exact characterization of these neurobiological processes is, however, essential for finding therapeutic strategies for affective or stress-related mental disorders. Here we provide compelling evidence of how brain networks respond to threat. We see a unified account of theta band driven network re-organization with transitions of certain brain regions within the threat, attention and salience circuits that interact during instructed threat processing. We based the timing of the TMS pulses to dMPFC on the dynamics of theta driven alterations, which return to baseline at 1000ms, as shown by inter-trial coherence in the experiment without TMS. The flexibility of the community structures and the characteristic of forming local clusters are dynamic network properties that predetermine the physiological responses to threat. Moreover, we see a time dependent decrease of threat induced network behavior (increased flexibility and clustering in the threat network), that can be causally modulated by an applied transcranial magnetic stimulation pulse over dMPFC 1000 ms after CS+ presentation. A test pulse applied over dMPFC at a time period not relevant for threat processing interfered significantly with ongoing network behavior and community restructuring.

Furthermore, the specific connection strengths between amygdala and insula or cortico-cortical identified prior to a subject’s involvement in threat processing predicted the threat response, as measured by heart rate increase or subjective threat rating. Similarly, the strength of these connections also predicted the dMPFC-TMS dependent modulation of network behavior and threat processing.

### Community reorganization through connectivity dynamics is required for threat processing

Our results show that baseline activity (before CS+ presentation) has clear functional separation in several communities ^47^. However, during threat processing, the modular topology of these communities was modified, leading to the formation of three new known networks: DAN, SN and FN. These networks and their core components have been described as participating in threat and emotional processing ^48^. Restructuring of resting to specific task-related networks therefore appears to be a primordial mechanism that mediates between perception of relevant inputs and appropriate subsequent higher-order processing. Our results further highlight the presence of parallel processing during threat, where the involved networks (FN, DAN, SN) may distribute separate aspects of high-order cognitive workflow in order to cope with the situation. Of note, this may differ from the processing of classical Pavlovian threat conditioning paradigms, since in the designed paradigm the subjects are previously instructed about the contingency between CS+ and US, so expect the threatening event ^49^. This necessitates recruitment of additional (attentional and control) resources parallel to those in needed during the conditioned threat responses.

Despite our results showing that the interplay between synchrony of oscillations and network architecture is a key factor to mediate and sustain efficient information transfer for longer time periods, an open question remains regarding the state-dependent dynamics of the network, especially its dependency on stimulus relevance. Here, the network flexibility emerges as a state-dependent parameter of the threat processing, evidenced by its increase in the three networks. The network parameter flexibility has been already shown to increase during tasks necessitating cognitive flexibility ^50^, suggesting that dynamic reconfiguration of brain networks boosts efficient threat processing.

### Inter-trial coherence as a substrate of threat processing

We found an increase of the theta inter-trial phase coherence in the frontal lobe and a simultaneous decrease in the inter-trial phase coherence for the occipital alpha, relative to stimuli presentation. This suggests that threat processing is not a purely autonomous response to stimulus presentation, but rather that it facilitates interactions between regions ^51^ and for specific temporal conditioned stimuli ^52^. Previous studies on memory function have shown that there is a relationship between brain response to external stressors and the phase of the synchronized oscillations ^3^, which can be prolonged by exciting a small number of neurons ^53^ that participate in such oscillatory behavior. Low-frequency oscillations, such as theta (4–7 Hz) and alpha (8–12 Hz) can be recorded in different specific anatomical regions and especially facilitate communication between hippocampus ^54^, amygdala ^55^ and prefrontal cortex. The specificity of attention in instructed threat studies suggests that these oscillations provide a temporal window for inter-regional communication ^56^. Intra-regional functional communication has been found for interactions involving the fronto-occipital circuit during directed attention to visual stimuli ^57,58^. It has also previously been shown that frontal theta phase consistency reflects coordination of information transfer between distant brain areas ^59,60^. On the contrary, the decrease in the inter-trial phase for the alpha oscillations in the occipital lobe could be related to alterations due to pre-visual threat processing. Earlier studies ^61,62^ have shown disruption in phase consistency over successive trials in the occipital lobe, which suggest that the inter-trial coherence of these oscillations drives the physiological response during instructed threat processing. Our data support this hypothesis and localize it differentially in both the frontal and occipital lobes. Here we use the dynamics of theta driven alterations for the application of TMS pulses to dMPFC, and apply TMS at a time relevant for threat processing (1000 ms) and a time point before this (80 ms). Using this approach, we were able to achieve different effects at the network behavior level. Specifically, a TMS pulse before active processing leads to a community independent increase of network flexibility and clustering without a preservation of inter-network interactions. A TMS pulse applied during a time point physiologically relevant for processing mirrors the network response and intercommunity information transfer.

### Modulation of information flow directionality is required for threat processing and its corresponding behavioral correlates

The results presented here further demonstrate causal network dynamics within the reconfigured networks during threat processing and with TMS pulses. The temporal changes associated with threat processing are predominantly mediated uni-directionally, as previously shown in low frequency oscillations of amygdala-hippocampus connections ^63^. More specifically, the network dynamics in the DAN take a more parietal to frontal uni-directional route during threat processing, which is in line with previous reports using Pavlovian conditioning paradigms ^64^. In the salience network, the connections also show strong uni-directional connectivity to threat processing, while TMS facilitates information processing in the regions composing this network. Moreover, a heightened response of the threat network to expected threat stimuli has been recently shown using startle responses ^65^. Accordingly, we aimed to test the hypothesis that a targeted modulation of a region of the threat processing network through TMS pulses can induce similar large-scale network dynamics ^66^ modulating reorganization and information flow among distant regions ^67^.

Behavioral responses have been shown to be good correlates of induced threat processing ^49,68^. Accordingly, significant increases for the CS+ condition in both threat ratings and heart rate were observed in both of our experiments. Significantly, however, neither the behavioral ratings nor quantitative heart rate responses were modulated by TMS pulses. It has been previously shown that local perturbations should not change the behavioral responses ^33,69^. The behavioral indicators correlated with specific connections in the three newly formed communities which involved cortical and cortical-subcortical routes. We were also able to validate the correlations with two experiments for some of the connections, demonstrating that the changes in the analyzed network dynamics that we observed at the time of stimulation were purely induced by the TMS.

### Conclusions

Our findings demonstrate that threat processing is related to changes in the brain’s modular architecture involving the dorsal attention, salience and threat networks. Changes in flexibility and local connectivity in these three networks is a prerequisite for threat processing and related to behavioral responses. The TMS modulated theta and alpha oscillations, changed the dynamics of network flexibility, and caused a decrease in the DAN and increase in the SN and FN. The observation of modulation at both local and global network levels for information directionality and network re-organization to threat processing and TMS stimulation suggest that these dynamical phenomena serve as adaptive mechanisms for efficient threat processing.

